# Steady-state distributions of nascent RNA for general initiation mechanisms

**DOI:** 10.1101/2022.03.30.486441

**Authors:** Juraj Szavits-Nossan, Ramon Grima

## Abstract

Fluctuations in the number of nascent RNA accurately reflect transcriptional activity. However, mathematical models predicting their distributions are difficult to solve analytically due to their non-Markovian nature stemming from transcriptional elongation. Here we circumvent this problem by deriving an exact relationship between the steady-state distribution of nascent RNA and the distribution of initiation times, which can be computed for any general initiation mechanism described by a set of first-order reactions. We test our theory using simulations and live cell imaging data.

---

Transcription in single cells occurs in bursts whose size and timing is random [1]. Intuitively, a burst originates from rapid mRNA transcription when a promoter briefly switches on. The experimental distribution of mature mRNA numbers can often be fitted using the exact solution of a simple Markov model of gene expression, called the telegraph model [2–5]. This model describes the promoter switching between two states of activity and inactivity, the production of mature mRNA occurring one molecule at a time whilst in the active state, and the degradation of mature mRNA via a first-order reaction. However, in recent years, doubts have arisen about the validity of this kinetic description, principally because mature (cellular) mRNA does not provide a direct readout of transcription [6, 7]—the fluctuations of mRNA numbers in the cell are strongly influenced by various post-transcriptional events such as splicing, nuclear export and DNA replication.

In order to circumvent such issues, it has been proposed that gene expression can be more accurately understood by studying nascent RNA fluctuations, i.e. variability in the number of RNA polymerase (RNAP) molecules that are actively involved in the elongation phase of transcription. By its very definition, this is a direct read-out of transcription. The numbers of actively transcribing RNAPs can be estimated directly from nascent single-cell sequencing methods [8] or more commonly using single molecule fluorescence in situ hybridization (sm-FISH) where intronic probes specifically label (non-spliced) nascent RNA [7, 9]. Fitting of these numbers using stochastic models that account for RNAP dynamics can help us more accurately understand which regulatory steps in transcription are tuned to achieve required mRNA expression levels [10, 11].

Real-time observation of transcription in *vivo* has revealed that initiation is a stochastic process, whereas elongation and termination are fairly deterministic [12]. Transcription can be thus modelled by a stochastic, multistep process describing initiation, followed by a deterministic, single-step reaction describing elongation and termination. Despite this simplification, models of this type are particularly difficult to solve because they are inherently non-Markovian. So far, only two such models have been solved analytically: one in which the promoter is always active [13, 14], and the other in which the promoter switches between two states of activity and inactivity (Fig. 1(b)) [6]. A few other, more realistic models have been studied, but only the first two moments of the nascent RNA distribution have been obtained analytically [15–17]. None of these models, however, account for the complex multistep process of initiation that has been elucidated by decades of biochemical research [18].

**FIG. 1.**
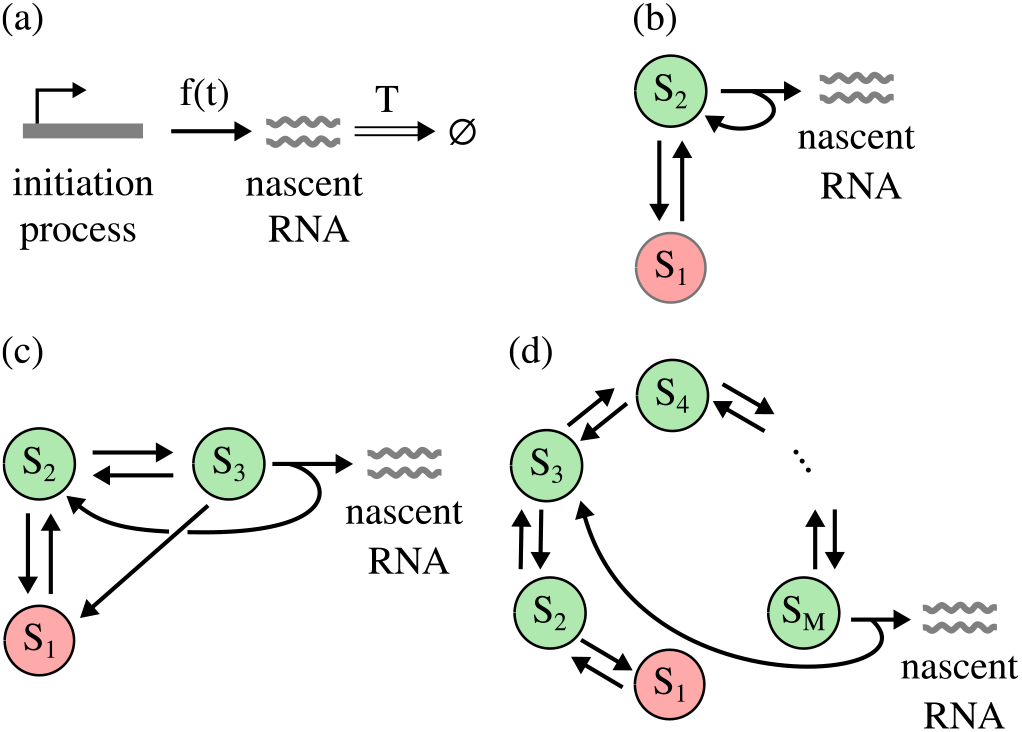
(a) Nascent RNA is produced by an initiation process at time intervals that are independent and identically distributed (iid) random variables with an arbitrary probability density function *f*(*t*). Elongation and termination are modelled deterministically, i.e. they take a fixed amount of time *T* to finish. (b)-(d) Examples of the initiation processes that can be studied by this framework, in the order of increasing complexity. *S*_1_, *S*_2_, etc are gene states. Red circles denote the off states. (b) The telegraph process [2]. (c) A three-state process that accounts for the binding of RNAP [19]. (d) A stepwise process of eukaryotic transcription that accounts for the binding of general transcription factors and RNAP [18, 20], and the promoter proximal pausing of RNAP in metazoans [21, 22].

In this Letter, we develop a general framework that allows us to find nascent RNA distribution for models with stochastic, multistep initiation, under the assumption of deterministic elongation and termination. The significance of this result is that it applies to any initiation process that produces nascent RNA at time intervals that are independent and identically distributed (iid) random variables with an arbitrary probability density function *f*(*t*) (Fig. 1(a)). This is true for any stochastic process described by first-order reactions with time-independent kinetic rates, some of which are shown in Fig. 1. We argue that while transcription initiation includes invariably many bimolecular steps, these are well approximated by pseudo first-order reactions because transcriptional machinery is generally abundant; furthermore, any variations of the rates are over timescales that are much longer than the elongation time and hence can be ignored [23]. For such processes, we solve a first passage time problem to get an analytic expression for *f*(*t*). The steadystate nascent RNA distribution is then computed using renewal theory, which generalizes the Poisson process by allowing for a non-exponential *f*(*t*). We verify our theory by stochastic simulations and using experimental results for the transcription kinetics in *Escherichia coli* [24].

## Waiting time distribution between successive nascent RNA production events

Transcription is divided into initiation, elongation and termination. Initiation is a complex multistep process which starts by the binding of an RNAP at the promoter, together with several transcription factors, and ends when the RNAP escapes the promoter and starts productive elongation. During elongation, RNAP traverses the gene and copies its sequence into a nascent RNA. The termination occurs at the gene end, leading to a mature RNA transcript.

We consider a stochastic initiation process consisting of *M* gene states labelled by *S*_1_,… *S_M_*. The gene switches between the states according to first-order reactions

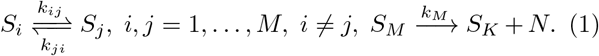

The last reaction describes production of nascent RNA (*N*), after which the initiation process starts again from the state *S_K_* (*K* takes a fixed value between 1 and *M*). Eq. 1 is a general description of the initiation process using first-order reactions. Particular cases are obtained by removing some reactions, which is equivalent to setting the rates of those reactions to zero.

We are interested in computing the probability density function (pdf) of the waiting time between successive events of nascent RNA production. This is a first passage time problem that can be solved by replacing the last reaction in Eq. 1 with *S_M_* → *A*, where *A* is an absorbing state, i.e. once the process reaches A, it stops. The pdf *f*(*t*) is then equal to *k_M_P_M_*(*t*), where *P_M_*(*t*) is the probability that the gene is in state *M* at time *t*, given that it was in state *K* at time 0, and *k_M_dt* is the probability that the transition *S_M_* → *A* occurs in the time interval [*t*, *t* + *dt*). The probability *P_M_*(*t*) can be found by solving the following master equation,

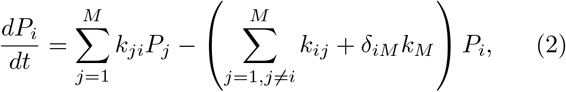

with the initial condition *P_i_*(0) = *δ_iK_*. The pdf *f*(*t*) is given by

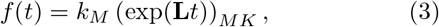

where the matrix elements of **L** read

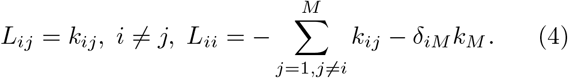

A central function in renewal theory is the Laplace transform of the pdf *f*(*t*), denoted by *f*\*(*s*). Formally,

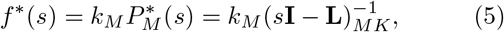

where 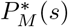 is the Laplace transform of *P_M_*(*t*) and **I** is the *M* × *M* identity matrix. Figs. 2(a)-(c) show *f*(*t*) for the initiation processes depicted in Fig. 1, computed from Eq. (5) and compared to the results of stochastic simulations. For the stepwise model of initiation in Fig. 1(d), *f*\*(*s*) can be computed analytically for any number of states M, because the matrix **L** is tridiagonal, for which the inverse is known explicitly [25]. The derivation of *f*\*(*s*) for all three models in Fig. 1 is presented in Supplemental Material [26].

**FIG. 2.**
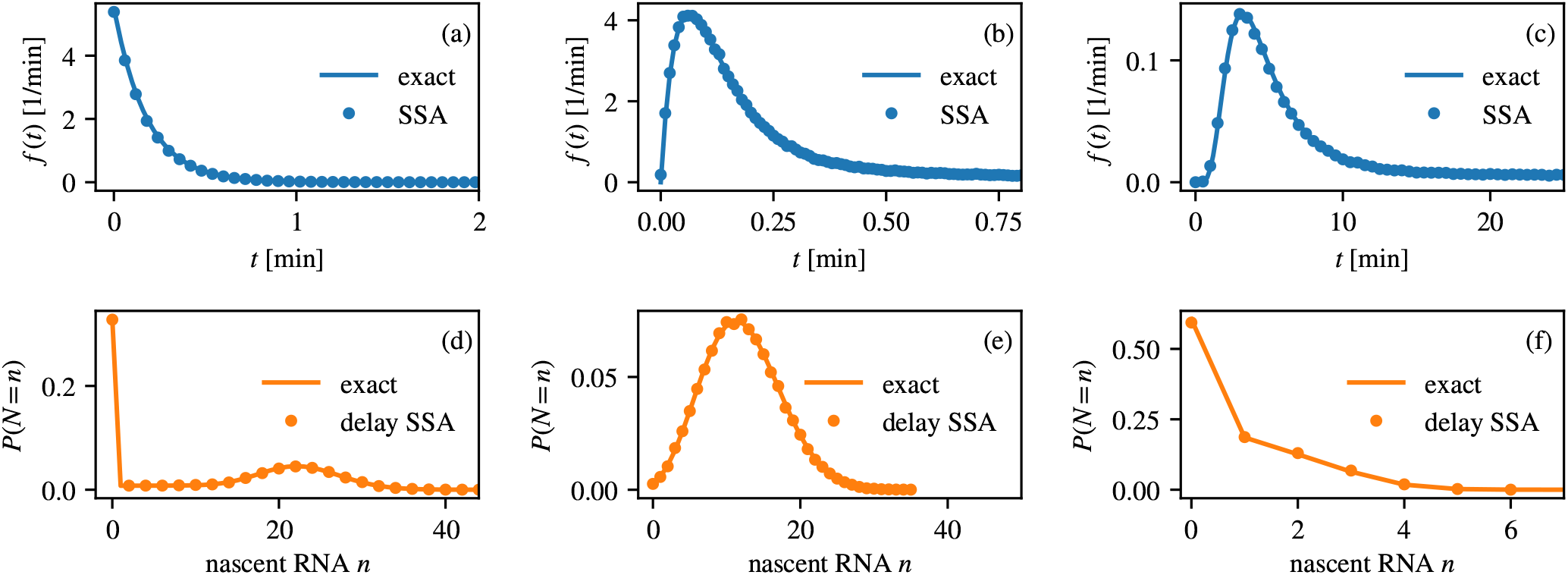
(a) to (c) Probability density function *f*(*t*) of the waiting time between successive nascent RNA production events for the initiation processes depicted in Fig. 1(b) to 1(d), respectively. Solid line is the theoretical prediction from Eq. (3), and points are from stochastic simulations (SSA). (b) The respective probability distribution of the nascent RNA, obtained from Eq. (9), and compared to stochastic simulations (delay SSA) performed using `DelaySSAToolkit.jl` package in Julia [26, 28]. Model parameters are listed in Supplemental Material [26].

## Distribution of nascent RNA in the steady state

Our approach for solving the general problem in Fig. 1(a) is based on recognizing that the number of nascent RNA production events that have occurred up to time *t*, denoted by *N_I_*(*t*), constitutes a renewal process [27]. A renewal process generalizes the Poisson process by allowing for non-exponentially distributed waiting times between successive events. As in the Poisson process, the waiting times in a renewal process are mutually independent. In transcription, this assumption is supported by experimental evidence [12].

We denote by *T_i_* the time of the *i*-th nascent RNA production event, measured from some reference time point *t* = 0. For simplicity, we assume that a nascent RNA production event occurred immediately before *t* = 0. If *t_i_* denotes the waiting time between *i*-th and (*i* – 1)-th nascent RNA production events, *t_i_* = *T_i_* – *T*_*i*–1_, where *T*_0_ = 0, then 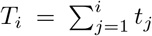. Since *t_j_* are iid random variables, the pdf of *T_i_* is the *i*-fold convolution of *f*, *k_i_*(*t*) = *f*^\**i*^(*t*).

The probability of *N_I_*(*t*) = *n* is equivalent to the probability of *T_n_* ≤ *t* < *T*_*n*+1_, which is given by

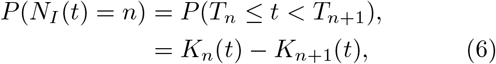

where 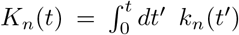 for *n* ≥ 1 is the cumulative distribution function of *T_n_*, and *K*_0_(*t*) = 1. Since the total time of elongation and termination is fixed, the number of nascent RNA actively engaged in transcription at time *t* is equal to

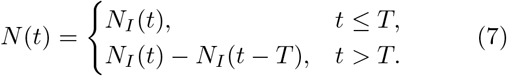

For *t* ≤ *T*, the probability *P*(*N*(*t*) = *n*) is given by Eq. (6). In order to find the probability distribution of *N*(*t*) for *t* > *T*, we first need to find the probability density function *f_t_0__*(*τ*) of the forward recurrence time *τ*, where *τ* is defined as the time until the next initiation event measured from a fixed time point *t*_0_ (we will later set *t*_0_ = *t* – *T*). In the steady state, *t*_0_ → ∞ and the expression for *f*_*t*_0__(*τ*) simplifies to

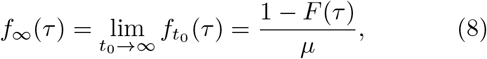

where 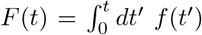 is the cumulative distribution function of the waiting times between successive nascent RNA production events, and *μ* is the mean waiting time [27]. Let *P*(*N* = *n*) denote the probability to find *n* nascent RNA on the gene in the steady state. Because nascent RNA production events are mutually independent, the Laplace transform *P*\*(*n, s*) of *P*(*N* = *n*) with respect to the elongation and termination time *T* is given by

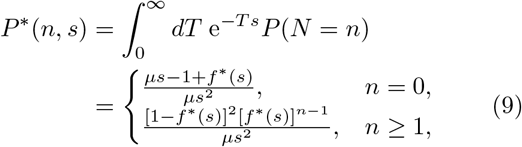

where we have used the fact that the Laplace transform of *f*_∞_(*τ*) is given by 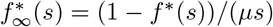.

Eq. (9) is our main result. It connects the Laplace transform of *P*(*N* = *n*) to the Laplace transform of *f*(*t*) for *all* initiation processes described by Eq. (2). For these processes, *f*\*(*s*) is a rational function of *s*, meaning that *P*(*N* = *n*) can be computed from Eq. (9) by partial fraction decomposition, for which many methods are available [29]. The moments of *P*(*N* = *n*) can be computed from the probability generating function 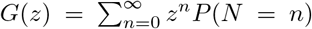, whose Laplace transform *G*\*(*z, s*) with respect to T is given by

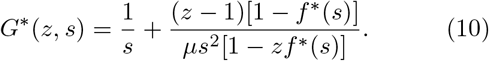

For example, the mean number of nascent RNA is equal to *T/μ*, and the variance is given by

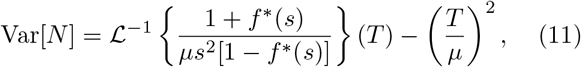

where 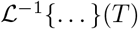 is the inverse Laplace transform of the expression in the curly brackets, evaluated at *T*.

Eq. (9) allows us to compute nascent RNA distribution analytically without using the complicated delay chemical master equation. In Supplemental Material, we demonstrate this for the Poisson process, the telegraph process and the fully irreversible process with equal forward rates *λ* [26]. In the last case, the waiting time distribution is an Erlang distribution, which is an important distribution that is found ubiquitously in cellular biology [30]. The nascent RNA distribution for this case reads

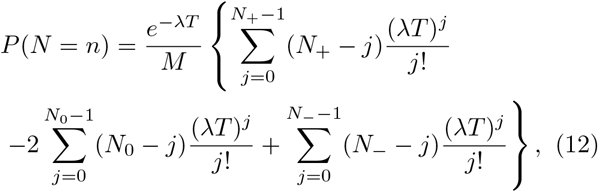

where *N*_+_ = *M*(*n* + 1), *N*_0_ = *Mn*, *N*_ = *M*(*n* – 1) [31].

Figs. 2(d)-(f) show the nascent RNA distribution for the initiation processes depicted in Fig. 1, computed from (9) and compared with the results of stochastic simulations. In particular, results in Figs. 2 (c) and (f) were obtained for a ten-state initiation process (*M* = 10) as a canonical model of eukaryotic transcription initiation [18, 20] using kinetic parameters collected from the literature [1, 32–39]. This model includes the on and off switching of the promoter, the binding and unbinding of six general transcription factors (IID, IIA, IIB, IIF, IIE and IIH) and RNAP, the unwinding of the doublestranded DNA, and the promotor proximal pausing of RNAP in metazoans [21, 22]. We assumed that the reinitiation starts from the on state (*K* = 2), whereas in Supplemental Material we consider the case in which transcription factors IID and IIA remain bound for a faster reinitiation (*K* = 4) [40, 41]. Sensitivity analysis for *K* = 2 revealed that the variability in nascent RNA is most sensitive to changes in the on and off rates, and the binding rate of TFIID, which indeed have been identified previously as rate-limiting steps in transcription initiation [19, 42].

## Comparison with experimental data

In Ref. [24], transcription kinetics of a target gene were followed in live *Escherichia coli* cells, one transcription event at a time. The target gene was controlled by the *tetA* promoter, which was induced (turned on) by anhydrotetracycline (aTc) at a concentration of 15 ng/ml at two temperatures, 24° C and 37° C. The waiting time distribution between successive mature RNA production events was measured experimentally and fitted to a hypoexponential distribution *f_fit_*(*t*) with rates *r*_1_, *r*_2_ and *r*_3_ (Fig. 3(a)). The Laplace transform of this distribution is given by

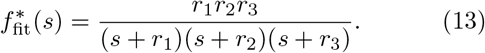

**FIG. 3.**
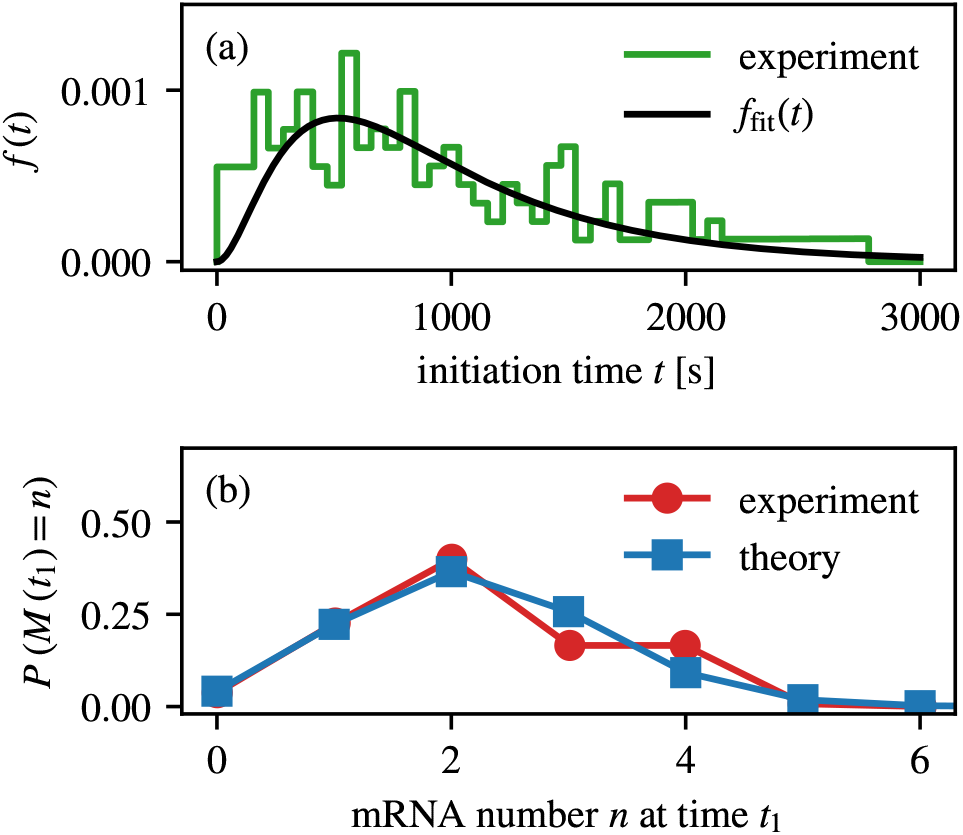
(a) Waiting time distribution between successive mature RNA production events measured in Ref. [24] at 24°C. Black line is a fit to the hypoexponential distribution with three parameters, *r*_1_ = 1/620 s^-1^, *r*_2_ = 1/240 s^-1^ and *r*_3_ = 1/115 s^-1^, obtained by inverting 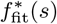 in Eq. (13). (b) Distribution of the fraction of cells with a given number of mRNA molecules measured 1 h following induction by aTc at 24°C, compared to the theoretical prediction obtained from Eq. (6) using 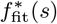 defined in Eq. (13).

Fig. 3(b) shows the distribution of the fraction of cells with a given number of mRNA, *M*(*t*), measured at *t* = *t*_1_ = 1 h after the induction by aTc at 24°C. This fraction is in a very good agreement with our theoretical prediction, which was computed as follows. The time of the full induction by aTC was estimated to 20 min [24]. We assumed that no transcription occurred before *t*_0_, and that the pdf of the waiting time until the first transcription event was *f*_fit_. Because mature RNA did not degrade during the experiment, we assumed that the number of mature RNA measured at *t*_1_, *M*(*t*_1_), equalled the number of individual transcription events between *t*_0_ and *t*_1_ – *T*, where *T* was the elongation and termination time. The latter was only tens of seconds, hence we approximated *t*_1_ – *T* by *t*_1_. The theoretical distribution *P*(*M*(*t*_1_) = *n*) was calculated from Eq. (6) by inverting the Laplace transform 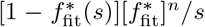, evaluated at *t*_1_ – *t*_0_ = 40 min. We stress that no fitting parameters were used other than those of *f*_fit_(*t*). The results at 37°C are also in good agreement with the experimental data (see Fig. S2(b) in Supplemental Material [26]). A very good agreement between the experiment and theory suggests that the knowledge of the waiting time distribution was sufficient to correctly predict the distribution of the accumulated mRNA.

## Conclusion

We have presented a general framework that allows us to predict the distribution of nascent RNA from the distribution of waiting times between successive nascent RNA production events, assuming the time of elongation and termination to be fixed. The significance of our solution is that it applies to any initiation mechanism that is modelled by first-order reactions with time-independent rates. The theory also allows for a model-free prediction of the nascent RNA distribution, provided the waiting time distribution is measured experimentally. Important directions for future work include assessing the importance of stochastic events in transcription elongation, such as RNAP collisions, ubiquitous pausing and premature termination, which have not been accounted for in this framework.

This work was supported by a Leverhulme Trust research award (RPG-2020-327).

