## Supplemental Material for "Steady-state distributions of nascent RNA for general initiation mechanisms"

Juraj Szavits-Nossan and Ramon Grima
*School of Biological Sciences, University of Edinburgh, Edinburgh EH9 3JH, United Kingdom*

**CONTENTS**

I. Supplemental derivations 1
A. Waiting time distributions for the initiation processes depicted in Fig. 1 in the main text 1
1. The telegraph process 1
2. Three-state process that accounts for the binding of RNAP 1
3. A stepwise process that accounts for the binding of general transcription factors and RNAP 2
B. Nascent RNA distributions for selected initiation processes 3
1. Poisson process 3
2. Telegraph process 4
3. Fully irreversible process with equal forward rates 6

II. Supplemental figures 8

III. Supplemental Tables 10

References 11

### I. SUPPLEMENTAL DERIVATIONS

#### A. Waiting time distributions for the initiation processes depicted in Fig. 1 in the main text

##### *1. The telegraph process*

This process has two gene states ($M = 2$). After initiation, the process returns to the state $S_2$ ($K = 2$). The conditional probability $P_i(t)$ that the gene is in state $S_i$ at time $t$, given that it was in state $S_2$ at time 0 and that no nascent RNA production has yet occurred, satisfies the following master equation

$$\begin{aligned} \frac{dP_1}{dt} &= k_{21}P_2 - k_{12}P_1, \\ \frac{dP_2}{dt} &= k_{12}P_1 - (k_2 + k_{21})P_2. \end{aligned} \tag{S1}$$

with the initial condition $P_i(0) = \delta_{i2}$. By taking the Laplace transform of both sides, we get

$$\begin{aligned} sP_1^* &= k_{21}P_2^* - k_{12}P_1^*, \\ sP_2^* - 1 &= k_{12}P_1^* - (k_2 + k_{21})P_2^*. \end{aligned} \tag{S2}$$

From here we get that the Laplace transform of the waiting time distribution $f(t)$ is given by

$$f^*(s) = \frac{k_2(s + k_{12})}{(s + k_{12})(s + k_{21} + k_2) - k_{21}k_{12}} = \frac{k_2(s + k_{12})}{s^2 + (k_{21} + k_{12} + k_2)s + k_{12}k_2}. \tag{S3}$$

##### *2. Three-state process that accounts for the binding of RNAP*

This process has three gene states ($M = 3$). After initiation, the process returns to the state $S_2$ ($K = 2$). The conditional probability $P_i(t)$ that the gene is in state $S_i$ at time $t$, given that it was in state $S_2$ at time 0 and that no nascent RNA production has yet occurred, satisfies the following master equation

$$\begin{aligned}\frac{dP_1}{dt} &= k_{31}P_3 + k_{21}P_2 - k_{12}P_1, \\ \frac{dP_2}{dt} &= k_{12}P_1 + k_{32}P_3 - (k_{21} + k_{23})P_2, \\ \frac{dP_3}{dt} &= k_{23}P_2 - (k_{31} + k_{32} + k_3)P_3,\end{aligned} \tag{S4}$$

with the initial condition $P_i(0) = \delta_{i2}$. By taking the Laplace transform of both sides, we get

$$\begin{aligned}sP_1^* &= k_{31}P_3^* + k_{21}P_2^* - k_{12}P_1^*, \\ sP_2^* - 1 &= k_{12}P_1^* + k_{32}P_3^* - (k_{21} + k_{23})P_2^*, \\ sP_3^* &= k_{23}P_2^* - (k_{31} + k_{32} + k_3)P_3^*.\end{aligned} \tag{S5}$$

From here we get that the Laplace transform of the waiting time distribution $f(t)$ is given by

$$f^*(s) = \frac{k_3k_{23}(s + k_{12})}{s^3 + (k_{12} + k_{21} + k_{23} + k_{32} + k_{31} + k_3)s^2 + [(k_{12} + k_{21})(k_{31} + k_{32} + k_3) + k_{23}(k_{12} + k_{31} + k_3)]s + k_{12}k_{23}k_3}. \tag{S6}$$

*3. A stepwise process that accounts for the binding of general transcription factors and RNAP*

This process has $M$ gene states. After initiation, the process returns to the state $S_K$. The conditional probability $P_i(t)$ that the gene is in state $S_i$ at time $t$, given that it was in state $S_K$ at time 0 and that no nascent RNA production has yet occurred, satisfies the following master equation

$$\begin{aligned}\frac{dP_1}{dt} &= k_{21}P_2 - k_{12}P_1, \\ \frac{dP_2}{dt} &= k_{12}P_1 + k_{32}P_3 - (k_{21} + k_{23})P_2, \\ &\vdots \\ \frac{dP_M}{dt} &= k_{M-1M}P_{M-1} - (k_{MM-1} + k_M)P_M,\end{aligned} \tag{S7}$$

with the initial condition $P_i(0) = \delta_{iK}$. We can write Eq. (S7) compactly as

$$\frac{dP}{dt} = \mathbf{A}P, \quad P(t) = \begin{bmatrix} P_1(t) \\ P_2(t) \\ \vdots \\ P_M(t) \end{bmatrix}, \quad \mathbf{A} = \begin{bmatrix} -e_1 & k_{21} & & & \\ k_{12} & -e_2 & k_{32} & & \\ & k_{23} & \ddots & \ddots & \\ & & \ddots & \ddots & k_{MM-1} \\ & & & k_{M-1M} & -e_M \end{bmatrix}, \tag{S8}$$

where $e_1 = k_{12}$, $e_i = k_{ii-1} + k_{ii+1}$ for $i = 2, \ldots, M-1$, $e_M = k_{MM-1} + k_M$, and the remaining matrix elements are zero. The solution to Eq. (S8) is

$$P(t) = e^{\mathbf{A}t}P(0), \tag{S9}$$

where $e^{\mathbf{A}t}$ is the matrix exponential. From here, we get the initiation time distribution as

$$f(t) = k_MP_M(t) = k_M\left(e^{\mathbf{A}t}\right)_{MK}, \tag{S10}$$

and from there, the Laplace transform of $f(t)$,

$$f^*(s) = k_M\left(s\mathbf{I} - \mathbf{A}\right)^{-1}_{MK}. \tag{S11}$$

Because $\mathbf{A}$ is a tridiagonal matrix, the inverse $(s\mathbf{I}-\mathbf{A})^{-1}$ can be found explicitly [1]. Following [1], we define the following recurrence relations,

$$z_0 = 1, \quad z_1 = s + e_1, \quad z_i = (s+e_i)z_{i-1} - k_{ii-1}k_{i-1i}z_{i-2}, \quad i = 2, \dots, M \tag{S12a}$$

$$y_{M+1} = 1, \quad y_M = s + e_M, \quad y_j = (s+e_j)y_{j+1} - k_{j+1j}k_{jj+1}y_{j+2}, \quad j = M-1\dots, 1. \tag{S12b}$$

The matrix element $(s\mathbf{I}-\mathbf{A})^{-1}_{MK}$ can be expressed as

$$(s\mathbf{I}-\mathbf{A})^{-1}_{M1} = \left(\prod_{n=1}^{M-1} k_{nn+1}\right) \frac{1}{y_2\left(s+e_1-k_{21}k_{12}\frac{y_3}{y_2}\right)}, \quad K=1 \tag{S13}$$

$$(s\mathbf{I}-\mathbf{A})^{-1}_{MK} = \left(\prod_{n=1}^{M-K} k_{nn+1}\right) \frac{1}{y_{K+1}\left(s+e_K-k_{KK-1}k_{K-1K}\frac{z_{K-2}}{z_{K-1}}-k_{K+1K}k_{KK+1}\frac{y_{K+2}}{y_{K+1}}\right)},$$
$$K = 2, \dots, M-1 \tag{S14}$$

$$(s\mathbf{I}-\mathbf{A})^{-1}_{MM} = \frac{1}{s+e_M-k_{MM-1}k_{M-1M}\frac{z_{M-2}}{z_{M-1}}}, \quad K = M, \tag{S15}$$

from which we get $f^*(s)$ by multiplying by $k_M$.

### B. Nascent RNA distributions for selected initiation processes

#### 1. *Poisson process*

The Poisson process is a special case of the stepwise model for $M=1$ and $K=1$,

$$S_1 \xrightarrow{k_1} S_1 + N. \tag{S16}$$

The waiting time distribution $f(t)$ is an exponential,

$$f(t) = k_1 e^{-k_1 t}, \quad f^*(s) = \frac{k_1}{s+k_1}, \tag{S17}$$

and the mean is given by $\mu = 1/k_1$. In this case (and in this case only), the forward recurrence time $\tau$ has the same exponential distribution as $f(t)$, $f_{t_0}(\tau) = k_1 e^{-k_1\tau}$, i.e. it does not depend on $t_0$, which is the memoryless property of the exponential distribution. For $t<T$, $P(N(t)=n)$ is a convolution of $f$ and $K_{n-1}-K_n$ evaluated at $t$,

$$\begin{aligned} P(N(t)=n) &= \int_0^t dt' f_{t-T}(t')[K_{n-1}(t-t') - K_n(t-t')] \\ &= \int_0^t dt' k_1 e^{-k_1 t'} \frac{1}{(n-1)!}(k_1(t-t'))^{n-1} e^{-k_1(t-t')} = \frac{(k_1 t)^n}{n!} e^{-k_1 t}, \quad t<T. \end{aligned} \tag{S18}$$

where we have used the fact that $K_n(t)$ is the Erlang distribution with the shape parameter $n$ and the rate parameter $k_1$,

$$K_n(t) = 1 - \sum_{m=0}^{n-1} \frac{(k_1 t)^m}{m!} e^{-k_1 t}. \tag{S19}$$

For $t>T$, $P(N(t)=n)$ is a convolution of $f_{t-T}$ and $K_{n-1}-K_n$ evaluated at $T$,

$$\begin{aligned} P(N(t)=n) &= \int_0^T dt' f_{t-T}(t')[K_{n-1}(t-t') - K_n(t-t')] \\ &= \int_0^T dt' k_1 e^{-k_1 t'} \frac{1}{(n-1)!}(k_1(t-t'))^{n-1} e^{-k_1(t-t')} = \frac{(k_1 T)^n}{n!} e^{-k_1 T}, \quad t>T. \end{aligned} \tag{S20}$$

Eq. (S20) is also the stationary distribution of the nascent RNA. Hence, the steady state is reached immediately after the first round of transcription elongation, i.e. at time $T$.

*2. Telegraph process*

The telegraph process is a special case the stepwise process for $M = 2$ and $K = 2$,

$$S_1 \underset{k_{21}}{\overset{k_{12}}{\rightleftharpoons}} S_2 \xrightarrow{k_2} S_2 + N. \tag{S21}$$

The Laplace transform of the waiting time distribution is given by Eq. (S3), and the mean $\mu$ is given by

$$\mu = -\left.\frac{df^*}{ds}\right|_{s=0} = \frac{k_{21}+k_{12}}{k_{12}k_2}. \tag{S22}$$

The Laplace transform of the nascent RNA distribution, $P^*(n,s)$, reads

$$P^*(n,s) = \begin{cases} \frac{\mu s - 1 + f^*(s)}{\mu s^2}, & n = 0 \\ \frac{[1-f^*(s)]^2[f^*(s)]^{n-1}}{\mu s^2}, & n \geq 1. \end{cases} \tag{S23}$$

We define $\lambda_1$ and $\lambda_2$ such that

$$(s+\lambda_1)(s+\lambda_2) = s^2 + s\,(k_{21}+k_{12}+k_2) + k_{12}k_2, \tag{S24}$$

from which it follows that

$$\lambda_{1,2} = \frac{k_{21}+k_{12}+k_2 \pm \sqrt{(k_{21}+k_{12}+k_2)^2 - 4k_{12}k_2}}{2}. \tag{S25}$$

Inserting Eq. (S3) into (S23) for $n = 0$, we get

$$P^*(0,s) = \frac{\mu s + \mu\lambda_1 + \mu\lambda_2 - 1}{\mu(s+\lambda_1)(s+\lambda_2)} = \frac{\lambda_2 - 1/\mu}{(\lambda_2-\lambda_1)(s+\lambda_1)} + \frac{\lambda_1 - 1/\mu}{(\lambda_1-\lambda_2)(s+\lambda_2)}. \tag{S26}$$

Using

$$\mathcal{L}^{-1}\left\{\frac{1}{(s+\alpha)^n}\right\}(T) = \frac{T^{n-1}}{(n-1)!}e^{-\alpha T}, \tag{S27}$$

we get

$$P(N=0) = \frac{\lambda_2 - 1/\mu}{\lambda_2 - \lambda_1}e^{-\lambda_1 T} + \frac{\lambda_1 - 1/\mu}{\lambda_1 - \lambda_2}e^{-\lambda_2 T}. \tag{S28}$$

For $n \geq 1$, we find

$$P^*(n,s) = \frac{1}{\mu}\left[\frac{s+k_{21}+k_{12}}{(s+\lambda_1)(s+\lambda_2)}\right]^2\left[\frac{(s+k_{12})k_2}{(s+\lambda_1)(s+\lambda_2)}\right]^{n-1} = \sum_{i=0}^{n}\frac{A_i}{(s+\lambda_1)^{n+1-i}} + \sum_{i=0}^{n}\frac{B_i}{(s+\lambda_2)^{n+1-i}}, \tag{S29}$$

where $A_i$ and $B_i$ are the coefficients in the partial fraction decomposition of $P^*(n,s)$,

$$A_i = \frac{1}{i!}\frac{d^i}{ds^i}\left[P^*(n,s)(s+\lambda_1)^{n+1}\right]\Big|_{s=-\lambda_1}, \quad i = 0,\ldots,n, \tag{S30}$$

$$B_i = \frac{1}{i!}\frac{d^i}{ds^i}\left[P^*(n,s)(s+\lambda_2)^{n+1}\right]\Big|_{s=-\lambda_2}, \quad i = 0,\ldots,n. \tag{S31}$$

From here, we get that the steady-state nascent RNA distribution for $n \geq 1$ is given by

$$\begin{aligned} P(N=n) &= \sum_{i=0}^{n}\frac{A_i}{(n-i)!}T^{n-i}e^{-\lambda_1 T} + \sum_{i=0}^{n}\frac{B_i}{(n-i)!}T^{n-i}e^{-\lambda_2 T} \\ &= \left(\frac{k_{12}}{k_{21}+k_{12}}\right)\frac{(k_2T)^n}{n!}\sum_{i=0}^{n}\binom{n}{i}\frac{\widetilde{A}_i e^{-\lambda_1 T} + \widetilde{B}_i e^{-\lambda_2 T}}{T^i}, \end{aligned} \tag{S32}$$

where $\widetilde{A}_i$ and $\widetilde{B}_i$ are given by

$$\widetilde{A}_i = i!A_i = \frac{d^i}{ds^i}\left[\frac{(s+k_{21}+k_{12})^2(s+k_{12})^{n-1}}{(s+\lambda_2)^{n+1}}\right]\Bigg|_{s=-\lambda_1}, \quad i=0,\dots,n, \tag{S33}$$

$$\widetilde{B}_i = i!B_i = \frac{d^i}{ds^i}\left[\frac{(s+k_{21}+k_{12})^2(s+k_{12})^{n-1}}{(s+\lambda_1)^{n+1}}\right]\Bigg|_{s=-\lambda_2}, \quad i=0,\dots,n. \tag{S34}$$

The remaining problem is to find the coefficients $\widetilde{A}_i$ and $\widetilde{B}_i$. To this end, we define auxiliary functions $u(s)$ and $v(s)$,

$$u(s) = \frac{(s+k_{12})^{n-1}}{(s+\lambda_2)^{n+1}}, \quad v(s) = \frac{(s+k_{12})^{n-1}}{(s+\lambda_1)^{n+1}}. \tag{S35}$$

The coefficient $\widetilde{A}_i$ can be written as,

$$\widetilde{A}_i = \begin{cases} (k_{21}+k_{12}-\lambda_1)^2u(-\lambda_1) & i=0, \\ (k_{21}+k_{12}-\lambda_1)^2u^{(1)}(-\lambda_1) + 2(k_{21}+k_{12}-\lambda_1)u(-\lambda_1) & i=1, \\ (k_{21}+k_{12}-\lambda_1)^2u^{(i)}(-\lambda_1) + 2i(k_{21}+k_{12}-\lambda_1)u^{(i-1)}(-\lambda_1) + i(i-1)u^{(i-2)}(-\lambda_1) & i=2,\dots,n. \end{cases}, \tag{S36}$$

where we have used the notation

$$u^{(i)}(-\lambda_1) = \frac{d^i}{ds^i}u(s)\Bigg|_{s=-\lambda_1}. \tag{S37}$$

By using the general Leibniz rule for differentiation, one can show that $u^{(i)}(-\lambda_1)$ can be written as

$$u^{(i)}(-\lambda_1) = (-1)^i(n+1)_i\,{}_2F_1\left(-i, 1-n, -i-n; \frac{\lambda_2-\lambda_1}{k_{12}-\lambda_1}\right)\frac{(k_{12}-\lambda_1)^{n-1}}{(\lambda_2-\lambda_1)^{n+1+i}}, \tag{S38}$$

where $(x)_i = x(x+1)\dots(x+i-1)$ is the rising factorial, and ${}_2F_1(a,b,c;z)$ is the hypergeometric function. The coefficient $\widetilde{B}_i$ can be obtained from $\widetilde{A}_i$ by replacing $u \leftrightarrow v$ and $\lambda_1 \leftrightarrow \lambda_2$,

$$\widetilde{B}_i = \begin{cases} (k_{21}+k_{12}-\lambda_2)^2v(-\lambda_2) & i=0, \\ (k_{21}+k_{12}-\lambda_2)^2v^{(1)}(-\lambda_2) + 2(k_{21}+k_{12}-\lambda_2)v(-\lambda_2) & i=1, \\ (k_{21}+k_{12}-\lambda_2)^2v^{(i)}(-\lambda_2) + 2i(k_{21}+k_{12}-\lambda_2)v^{(i-1)}(-\lambda_2) + i(i-1)v^{(i-2)}(-\lambda_2) & i=2,\dots,n. \end{cases}, \tag{S39}$$

where

$$v^{(i)}(-\lambda_2) = \frac{d^i}{ds^i}v(s)\Bigg|_{s=-\lambda_2} = (-1)^i(n+1)_i\,{}_2F_1\left(-i, 1-n, -i-n; \frac{\lambda_1-\lambda_2}{k_{12}-\lambda_2}\right)\frac{(k_{12}-\lambda_2)^{n-1}}{(\lambda_1-\lambda_2)^{n+1+i}}. \tag{S40}$$

By inserting Eq. (S3) into Eq. (9) in the main text, we get the following expression for the Laplace transform of the probability generating function,

$$G^*(z,s) = \frac{1}{s} + \frac{u(s+k_{21}+k_{12})}{\mu s[s^2+s(k_{21}+k_{12}-k_2u)-k_{12}k_2u]}, \tag{S41}$$

where $u = z-1$. Next, we introduce $\lambda_1(u)$ and $\lambda_2(u)$,

$$\lambda_{1,2}(u) = \frac{k_{21}+k_{12}-k_2u \pm \sqrt{(k_{21}+k_{12}-k_2u)^2+4k_{12}k_2u}}{2}, \tag{S42}$$

such that

$$s^2+s(k_{21}+k_{12}-k_2u)-k_{12}k_2u = [s+\lambda_1(u)]\,[s+\lambda_2(u)]\,. \tag{S43}$$

From there, we get

$$G^*(u,s) = \frac{1}{s} + \frac{u(s+k_{21}+k_{12})}{\mu s\,[s+\lambda_1(u)]\,[s+\lambda_2(u)]}, \tag{S44}$$

which can be inverted using the partial fraction decomposition. The final result for $G(u)$ is

$$G(u) = \frac{e^{-\lambda_1(u)T}}{2(k_1+k_{21})\sqrt{\Delta(u)}} \Big\{ (k_1+k_{21}) \left[\sqrt{\Delta(u)} - (k_1+k_{21})\right] - (k_1-k_{21})k_2 u \\ + (k_1+k_{21}) \left[\sqrt{\Delta(u)} + (k_1+k_{21})\right] e^{\sqrt{\Delta(u)}T} + (k_1-k_{21})k_2 u\, e^{\sqrt{\Delta(u)}T} \Big\}, \tag{S45}$$

where

$$\Delta(u) = (k_{21}+k_{12}-k_2 u)^2 + 4k_{12}k_2 u. \tag{S46}$$

The expression for $G(u)$ is the same as in Supplemental Material of Ref. [2], Eq. (1.15). There it was shown that the nascent RNA distribution, obtained from $G(u)$, matches the one obtained in Ref. [3].

#### 3. Fully irreversible process with equal forward rates

The reaction scheme for this initiation process is given by

$$S_1 \xrightarrow{\lambda} S_2 \xrightarrow{\lambda} \dots \xrightarrow{\lambda} S_M \xrightarrow{\lambda} S_1 + N. \tag{S47}$$

The initiation time distribution $f(t)$ is an Erlang distribution,

$$f(t) = \frac{\lambda^M t^{M-1}}{(M-1)!} e^{-\lambda t}, \quad f^*(s) = \frac{\lambda^M}{(s+\lambda)^M}, \tag{S48}$$

and the mean is given by $\mu = M/\lambda$. For $n = 0$, the inverse Laplace transform $P^*(0,s)$ is given by

$$P^*(0,s) = \frac{(Ms-\lambda)(s+\lambda)^M + \lambda^{M+1}}{Ms^2} \frac{1}{(s+\lambda)^M} = \sum_{i=0}^{M-1} \frac{A_i}{(s+\lambda)^{M-i}}, \tag{S49}$$

where the coefficient $A_i$ is given by

$$A_i = \frac{1}{i!} \frac{d^i}{ds^i} \left[ \frac{(s+\lambda)^M}{s} - \frac{\lambda}{M} \frac{(s+\lambda)^M}{s^2} + \frac{\lambda^{M+1}}{Ms^2} \right]\Bigg|_{s=-\lambda}, \quad i = 0, \dots, M-1 \tag{S50}$$

Because $i < M$, only the last term in the square brackets in Eq. (S50) contributes to $A_i$. The final result is

$$P(N=0) = \frac{e^{-\lambda T}}{M} \sum_{i=0}^{M-1} (M-j) \frac{(\lambda T)^i}{i!}. \tag{S51}$$

For $n \geq 1$, we get

$$P^*(n,s) = \frac{\lambda^{M(n-1)+1}}{M} \left[ \frac{(s+\lambda)^M - \lambda^M}{s} \right]^2 \frac{1}{(s+\lambda)^{M(n+1)}} = \sum_{i=0}^{M(n+1)-1} \frac{B_i}{(s+\lambda)^{M(n+1)-i}}, \tag{S52}$$

where $B_i$ is given by

$$B_i = \frac{1}{i!} \frac{d^i}{ds^i} \left[ P^*(n,s)(s+\lambda)^{M(n+1)} \right]\Big|_{s=-\lambda} = \frac{\lambda^{M(n-1)+1}}{Mi!} \frac{d^i}{ds^i} \left[ \frac{(s+\lambda)^{2M}}{s^2} - \frac{2\lambda^M (s+\lambda)^M}{s^2} + \frac{\lambda^{2M}}{s^2} \right]\Bigg|_{s=-\lambda}. \tag{S53}$$

For the derivatives of each term in the square brackets we get,

$$\frac{d^i}{ds^i} \frac{(s+\lambda)^{2M}}{s^2}\Bigg|_{s=-\lambda} = \begin{cases} i!(i-2M+1)\lambda^{2M-2-i}, & i \geq 2M \\ 0 & \text{otherwise} \end{cases}, \tag{S54}$$

$$\frac{d^i}{ds^i} \frac{2\lambda^M (s+\lambda)^M}{s^2}\Bigg|_{s=-\lambda} = \begin{cases} 2(i!)(i-M+1)\lambda^{2M-2-i}, & i \geq M \\ 0 & \text{otherwise} \end{cases}, \tag{S55}$$

$$\frac{d^i}{ds^i}\frac{\lambda^{2M}}{s^2}\bigg|_{s=-\lambda} = \begin{cases} (i+1)!\lambda^{2M-2-i}, & i \geq 0 \\ 0 & \text{otherwise} \end{cases}. \tag{S56}$$

By inserting these terms into the expression for $P^*(n,s)$ and taking the inverse Laplace transform, we get the following expression for the nascent RNA distribution,

$$P(N=n) = \frac{e^{-\lambda T}}{M}\left\{ \sum_{i=0}^{M(n-1)-1} [M(n-1)-j]\frac{(\lambda T)^i}{i!} - 2\sum_{i=0}^{Mn-1}(Mn-j)\frac{(\lambda T)^i}{i!} \right.$$
$$\left. + \sum_{i=0}^{M(n+1)-1} [M(n+1)-j]\frac{(\lambda T)^i}{i!} \right\}. \tag{S57}$$

Here, we have used a convention according to which a sum in which the upper bound is lower than the lower bound is equal to zero.

### II. SUPPLEMENTAL FIGURES

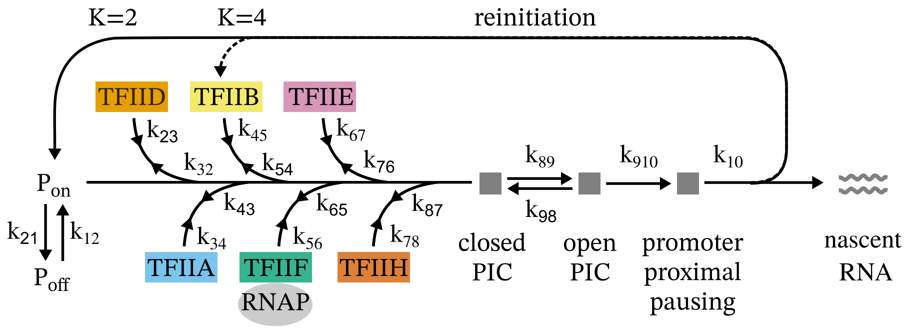


FIG. S1. A stepwise model of eukaryotic transcription initiation [4, 5] with ten gene states ($M = 10$). $\mathrm{P_{on}}$ ($S_1$) and $\mathrm{P_{off}}$ ($S_2$) are the on and off states of the promoter, respectively. General transcription factors and RNA polymerase bind the promoter in the following order: TFIID ($S_3$), TFIIA ($S_4$), TFIIB ($S_5$), TFIIF and RNAP ($S_6$), TFIIE ($S_7$) and TFIIH, resulting in the closed preinitiation complex (PIC, $S_8$). The TFIIH unwinds the promoter DNA, creating an open PIC ($S_9$) that begins the elongation. In metazoans, the elongating RNA polymerase pauses shortly after the initiation ($S_{10}$) [6, 7]. The RNAP is eventually released into productive elongation, clearing the promoter for the next round of initiation. We assumed that the reinitiation starts from the on state ($K = 2$). Alternatively, transcription factors IID and IIA may remain bound until a new round of initiation [8, 9]—this scenario corresponds to $K = 4$ (dashed line).

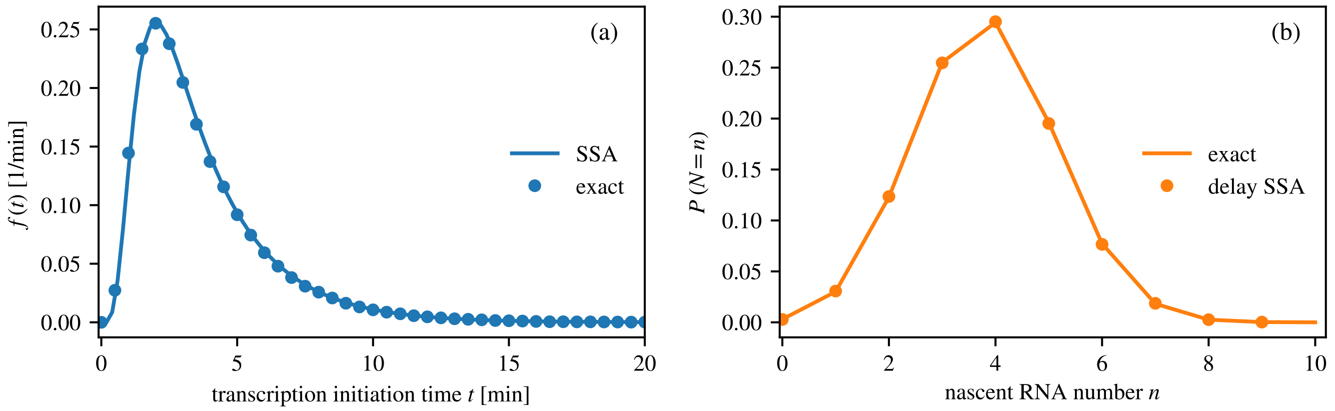


FIG. S2. (a) Probability density function $f(t)$ of the waiting time between successive nascent RNA production events for the stepwise model of eukaryotic transcription initiation illustrated in Fig. S1 with reinitiation from the promoter state with TFIID and TFIIA bound ($K = 4$). Solid line is the theoretical prediction from Eq. (3) in the main text, and points are from stochastic simulations (SSA). (b) Probability distribution of the nascent RNA, obtained from Eq. (9) in the main text, and compared to stochastic simulations (delay SSA) performed using `DelaySSAToolkit.jl` package in Julia [10, 11]. Parameters of the model are presented in Table S4.

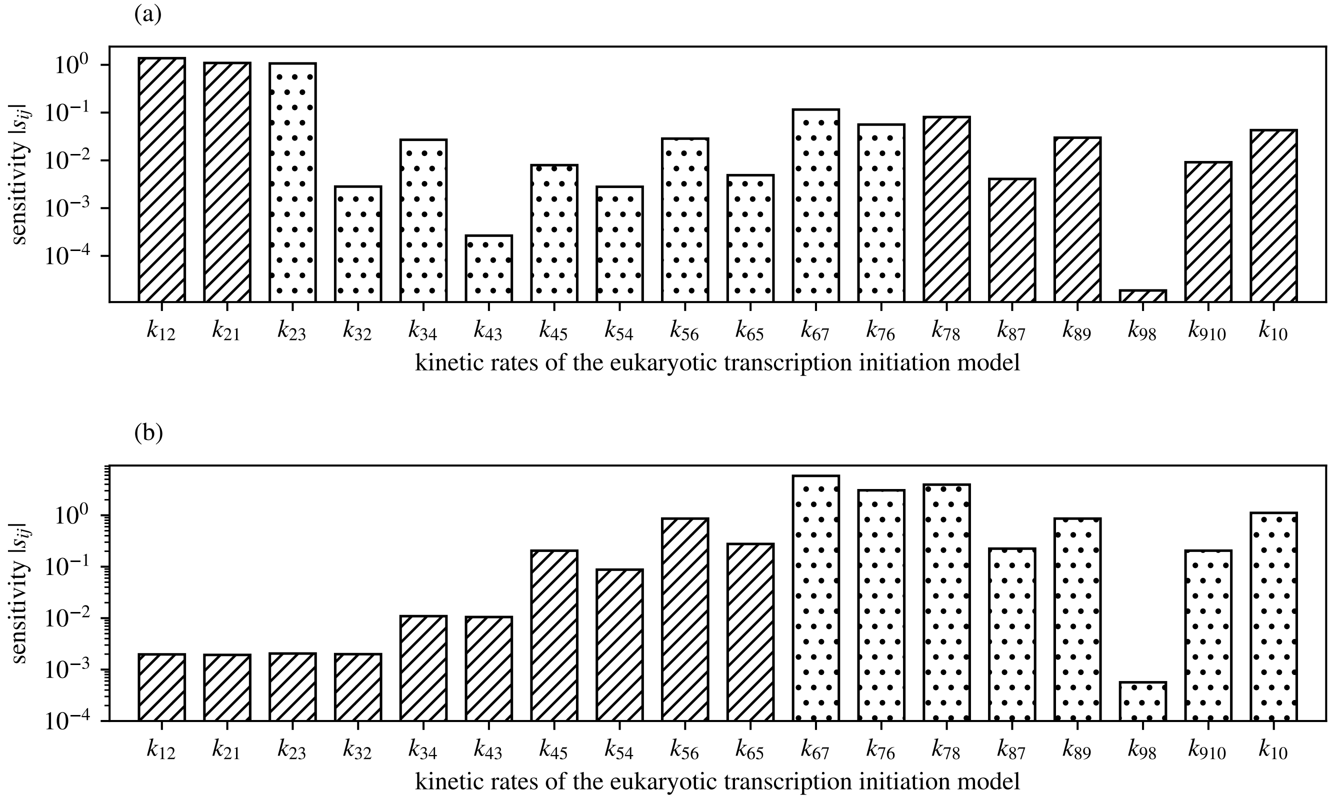


FIG. S3. Absolute value of the local sensitivity coefficient $|s_{ij}|$ of the Fano factor $\mathrm{FF}_N = \mathrm{Var}[N]/\langle N\rangle$ of the nascent RNA distribution, defined as $s_{ij} = (k_{ij}/\mathrm{FF}_n)\partial \mathrm{FF}_N/\partial k_{ij}$. Rates are labelled according to Fig. S1. Bars with diagonal lines have $s_{ij} < 0$, whereas bars with dots have $s_{ij} > 0$. (a) $K = 2$. (b) $K = 4$.

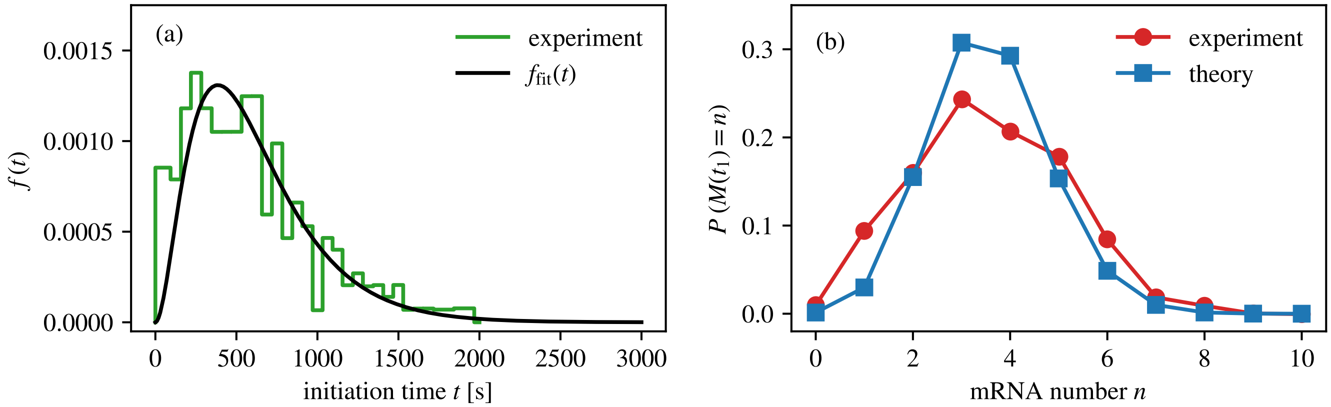


FIG. S4. (a) Distribution of the waiting times between successive mature RNA production events, measured in Ref. [12] at 37°C. Black line is a fit to the hypoexponential distribution with three parameters, $r_1 = 1/245\ \mathrm{s}^{-1}$, $r_2 = 1/254\ \mathrm{s}^{-1}$ and $r_3 = 1/109\ \mathrm{s}^{-1}$, obtained by inverting $f^*_{\mathrm{fit}}(s)$ in Eq. (13) in the main text. (b) Distribution of the fraction of cells with a given number of mRNA molecules measured 1 h following induction by aTc at 37°C, compared to the theoretical prediction obtained from Eq. (6) using $f^*_{\mathrm{fit}}(s)$ defined in Eq. (13) in the main text.

### III. SUPPLEMENTAL TABLES

TABLE S1. List of files that were used to solve transcription models of Fig. 1 in the main text. The files are available at https://github.com/jszavits/nascentRNA.

| Filename | Description |
|---|---|
| `model-1-SSA.ipynb` | Jupyter notebook file with Julia code; using stochastic simulations, it computes the waiting time distribution $f(t)$ for the initiation process depicted in Fig. 1(b) of the main text, and the corresponding nascent RNA distribution in the steady state |
| `model-1-exact.nb` | Mathematica notebook file; it computes the waiting time distribution $f(t)$ and the nascent RNA distribution using inverse Laplace transform |
| `model-2-SSA.ipynb` | Jupyter notebook file with Julia code; using stochastic simulations, it computes the waiting time distribution $f(t)$ for the initiation process depicted in Fig. 1(c) of the main text, and the corresponding nascent RNA distribution in the steady state |
| `model-2-exact.nb` | Mathematica notebook file; it computes the waiting time distribution $f(t)$ and the nascent RNA distribution using inverse Laplace transform |
| `model-3-SSA.ipynb` | Jupyter notebook file with Julia code; using stochastic simulations, it computes the waiting time distribution $f(t)$ for the initiation process depicted in Fig. 1(d) of the main text, and the corresponding nascent RNA distribution in the steady state |
| `model-3-exact.nb` | Mathematica notebook file; it computes the waiting time distribution $f(t)$ and the nascent RNA distribution using inverse Laplace transform |

TABLE S2. Parameter selection for the transcription model with the telegraph initiation process portrayed in Fig. 1(b) in the main text.

| Parameter | Description | Value | Reference |
|---|---|---|---|
| $k_{12}$ | promoter on rate | 0.06 $\text{min}^{-1}$ | - |
| $k_{21}$ | promoter off rate | 0.042 $\text{min}^{-1}$ | - |
| $k_2$ | initiation rate | 5.52 $\text{min}^{-1}$ | - |
| $L$ | length of the gene | 10 kb | - |
| $v$ | RNA polymerase speed | 2.4 kb/min | [7] |
| $T$ | elongation time | 4.167 min | $L/v$ |

TABLE S3. Parameter selection for the transcription model with the three-state initiation process portrayed in Fig. 1(c) in the main text.

| Parameter | Description | Value | Reference |
|---|---|---|---|
| $k_{12}$ | promoter on rate | 1.92 $\text{min}^{-1}$ | - |
| $k_{21}$ | promoter off rate | 1.92 $\text{min}^{-1}$ | - |
| $k_{23}$ | RNAP binding rate | 9.6 $\text{min}^{-1}$ | - |
| $k_{32}$ | RNAP unbinding rate | 0.96 $\text{min}^{-1}$ | - |
| $k_{31}$ | RNAP unbinding and promoter off rate | 1.92 $\text{min}^{-1}$ | - |
| $k_3$ | initiation rate | 19.2 $\text{min}^{-1}$ | - |
| $L$ | length of the gene | 10 kb | - |
| $v$ | RNA polymerase speed | 2.4 kb/min | [7] |
| $T$ | elongation time | 4.167 min | $L/v$ |

TABLE S4. Parameter selection for the model of eukaryotic transcription with ten gene states portrayed in Fig. 1(d) in the main text and also in Fig. S1. The initial values for $k_{23}, \ldots, k_{10}$ and $k_{32}, \ldots, k_{98}$, assumed to be representative of eukaryotes, were collected from the literature. The values of $k_{12}$ and $k_{21}$ were matched to the on and off rates inferred from the transcription kinetics of PLEC gene promoter in mouse fibroblast cells [13]. For this promoter, the mean time it took to initiate from the on state, assuming no return to the off state, was reported to be 5.2 min [13]. The initial values of $k_{54}, k_{56}, k_{89}$ and $k_{10}$ were adjusted to match this value to the mean initiation time obtained from $f(t)$ in Eq. (S10) after setting $k_{21} = 0$ and $K = 2$. The last three parameters describe the elongation stage. The elongation time $T$ was obtained by dividing the gene length of PLEC gene by the RNA polymerase II speed of 4.3 kb/min [14]. The values in bold have been changed from their initial values found in the literature to match the mean initiation time of 5.2 min from the on state, measured for the PLEC gene promoter in mouse [13].

| Parameter | Description | Initial Value | Final Value | Reference |
|---|---|---|---|---|
| $k_{12}$ | promoter on rate | $0.04$ min$^{-1}$ | $0.04$ min$^{-1}$ | [13] |
| $k_{21}$ | promoter off rate | $0.53$ min$^{-1}$ | $0.53$ min$^{-1}$ | [13] |
| $k_{23}$ | TFIID binding rate | $1/1.1$ min$^{-1}$ | $1/1.1$ min$^{-1}$ | [15] |
| $k_{32}$ | TFIID unbinding rate | $1/130$ min$^{-1}$ | $1/130$ min$^{-1}$ | [15] |
| $k_{34}$ | TFIIA binding rate | $1/20$ s$^{-1}$ | $1/20$ s$^{-1}$ | [16] |
| $k_{43}$ | TFIIA unbinding rate | $1/8$ min$^{-1}$ | $1/8$ min$^{-1}$ | [17] |
| $k_{45}$ | TFIIB binding rate | $1/3.2$ s$^{-1}$ | $1/3.2$ s$^{-1}$ | [16] |
| $k_{54}$ | TFIIB unbinding rate | $1/1.5$ s$^{-1}$ | $\mathbf{1/30}$ s$^{-1}$ | [16] |
| $k_{56}$ | TFIIF and RNAP binding rate | $2.3 \cdot 10^{-3}$ s$^{-1}$ | $\mathbf{6.9 \cdot 10^{-2}}$ s$^{-1}$ | [18] |
| $k_{65}$ | TFIIF and RNAP unbinding rate | $3 \cdot 10^{-3}$ s$^{-1}$ | $3 \cdot 10^{-3}$ s$^{-1}$ | [18] |
| $k_{67}$ | TFIIE binding rate | $1.7 \cdot 10^{-2}$ s$^{-1}$ | $1.7 \cdot 10^{-2}$ s$^{-1}$ | [19] |
| $k_{76}$ | TFIIE unbinding rate | $5 \cdot 10^{-2}$ s$^{-1}$ | $5 \cdot 10^{-2}$ s$^{-1}$ | [19] |
| $k_{78}$ | TFIIH binding rate | $5 \cdot 10^{-2}$ s$^{-1}$ | $5 \cdot 10^{-2}$ s$^{-1}$ | [19] |
| $k_{87}$ | TFIIH unbinding rate | $1/5$ min$^{-1}$ | $1/5$ min$^{-1}$ | [19] |
| $k_{89}$ | closed to open PIC | $1.9 \cdot 10^{-3}$ s$^{-1}$ | $\mathbf{5.7 \cdot 10^{-2}}$ s$^{-1}$ | [20] |
| $k_{98}$ | open to closed PIC | $1.1 \cdot 10^{-4}$ s$^{-1}$ | $1.1 \cdot 10^{-4}$ s$^{-1}$ | [20] |
| $k_{910}$ | open PIC to elongation | $0.17$ s$^{-1}$ | $0.17$ s$^{-1}$ | [20] |
| $k_{10}$ | escape rate from the promoter proximal pausing | $1/5$ min$^{-1}$ | $\mathbf{1/0.5}$ min$^{-1}$ | [21] |
| $L$ | length of PLEC gene in mouse | 60.404 kb | 60.404 kb | Gene ID: 18810 |
| $v$ | RNA polymerase speed | 4.3 kb/min | 4.3 kb/min | [14] |
| $T$ | elongation time | 14 min | 14 min | $L/v$ |

[11] See Supplemental Material at [URL will be inserted by publisher] for additional derivations, figures, tables with model parameters, and computer programs for solving the transcription models from Fig. 1.
